# Children with autism observe social interactions in an idiosyncratic manner

**DOI:** 10.1101/720706

**Authors:** Inbar Avni, Gal Meiri, Asif Bar-Sinai, Doron Reboh, Liora Manelis, Hagit Flusser, Analya Michaelovski, Idan Menashe, Ilan Dinstein

## Abstract

Previous eye tracking studies have reported that children with autism spectrum disorders (ASD) fixate less on faces in comparison to controls. To properly understand social interactions, however, children must gaze not only at faces, but also at actions, gestures, body movements, contextual details, and objects, thereby creating specific gaze patterns when observing specific interactions. We presented three different movies of social interactions to 111 children (71 with ASD) who watched each of the movies twice. Typically developing children viewed the movies in a remarkably predictable and reproducible manner, exhibiting gaze patterns that were similar to the mean gaze pattern of other controls, with strong correlations across individuals (inter-subject correlations) and across movie presentations (intra-subject correlations). In contrast, children with ASD exhibited significantly more variable/idiosyncratic gaze patterns that differed from the mean gaze pattern of controls and were weakly correlated across individuals and presentations. Most importantly, quantification of gaze idiosyncrasy in individual children, enabled separation of ASD and control children with higher sensitivity and specificity than traditional measures such as time gazing at faces. Individual magnitudes of gaze idiosyncrasy were also significantly correlated with ASD severity and significantly correlated across movies and movie presentations, demonstrating their clinical sensitivity and reliability. These results suggest that gaze idiosyncrasy is a potent behavioral abnormality that characterizes many children with ASD and may contribute to their impaired social development. Quantification of gaze idiosyncrasy in individual children may aid in assessing their ASD severity over time and in response to treatments.

**Lay Summary:** Typically developing children watch movies of social interactions in a reliable and predictable manner, attending faces, gestures, body movements, and objects that are relevant to the social interaction and its narrative. Here, we demonstrate that children with ASD watch such movies with significantly more variable/idiosyncratic gaze patterns that differ across individuals and across movie presentations. We demonstrate that quantifying this variability is a very potent way of identifying children with ASD and determining the severity of their social ASD symptoms.

## Introduction

Throughout life we continuously select our visual input by actively controlling gaze position (Henderson, 2003; Schroeder, Wilson, Radman, Scharfman, & Lakatos, 2010). Gaze behavior, therefore, governs the exposure that a child has to social stimuli and their ability to learn social skills through experience dependent plasticity (Hensch, 2005). When observing social interactions, human gaze behavior is often attracted to faces, which contain important social information regarding the intentions, feelings, and goals of others (Baron-Cohen, Wheelwright, & Jolliffe, 1997; Ekman & Friesen, 1971). This preference for faces is evident already in infants during their first months of life (Batki, Baron-Cohen, Wheelwright, Connellan, & Ahluwalia, 2000; Frank, Vul, & Johnson, 2009; Johnson, 2005). However, a preference for faces is not the only factor governing gaze behavior. Additional factors that are apparent in typically developing toddlers include an attraction to visually salient stimuli (Helo, van Ommen, Pannasch, Danteny-Dordoigne, & Rämä, 2017), biological motion (Fox & McDaniel, 1982; Simion, Regolin, & Bulf, 2008), following the gaze of others (related to joint attention) (Argyle & Cook, 1976; Emery, 2000), and observing the targets of others actions (related to theory of mind) (Flanagan & Johansson, 2003; Oniski & Baillargeon, 2005). Together, these factors and others create convergence and correlation in the gaze patterns of typically developing children (Constantino et al., 2017; Franchak, Heeger, Hasson, & Adolph, 2016) and adults (Shepherd, Steckenfinger, Hasson, & Ghazanfar, 2010; Wang, Freeman, Merriam, Hasson, & Heeger, 2012) during the observation of movies containing social interactions.

It has been proposed that gaze abnormalities in children with ASD may reduce their early exposure to social information and impair their ability of learn basic social skills (Klin, Shultz, & Jones, 2015). Indeed, a common behavioral symptoms of ASD is reduced eye contact (Senju & Johnson, 2009; Tanaka & Sung, 2016) and previous eye tracking studies have reported that children with ASD exhibit weaker gaze preferences for people (Moore et al., 2018; Pierce et al., 2016), faces (Chawarska, Macari, & Shic, 2012; Chita-Tegmark, 2016; Constantino et al., 2017; W Jones, Carr, & Klin, 2008; Warren Jones & Klin, 2013; Papagiannopoulou, Chitty, Hermens, Hickie, & Lagopoulos, 2014; Riby & Hancock, 2009; Rice, Moriuchi, Jones, & Klin, 2012; Q. Wang, Campbell, Macari, Chawarska, & Shic, 2018), biological motion (Falck-Ytter, Rehnberg, & Bölte, 2013; Klin, Lin, Gorrindo, Ramsay, & Jones, 2009), and following the gazes of others (Bedford et al., 2012). These studies have suggested that quantifying gaze behavior with eye-tracking may be a potent technique for estimating the initial severity of social symptoms in ASD and sensitively tracking their change over time or in response to treatments (Frazier et al., 2018; Sasson & Elison, 2012).

The studies described above have quantified gaze abnormalities in children with ASD using two types of measures. The first estimates the relative amount of time that children gaze at manually defined regions of interest (ROIs) within each frame of the movie. ROIs typically include the face, eyes, mouth, body, objects, or other items of potential interest (e.g., an object that is being manipulated) (Chawarska et al., 2012; W Jones et al., 2008; Warren Jones & Klin, 2013). The second estimates the relative amount of time that children gaze at each side of a split screen that contains two different stimuli (e.g., children exercising on one side and geometrical shapes on the other (Moore et al., 2018; Pierce et al., 2016)). Both of these measures estimate gaze abnormalities using summary statistics that quantify the total amount of time that a child gazes at a particular stimulus, regardless of when they gazed at it (i.e., ignoring the temporal gaze pattern).

An alternative approach is to compare the actual moment-by-moment gaze patterns of ASD and typically developing children during observation of movies, given that movies are known to create strong correlation in the gaze patterns of neuro-typical individuals (Franchak et al., 2016; Shepherd et al., 2010; H. Wang et al., 2012). With this in mind, two studies have used multidimensional scaling (Nakano et al., 2010) or cohesion (Wang et al., 2018) measures to demonstrate that gaze patterns of individuals with ASD are more variable than those of controls. To date, the utility of different measures for identifying ASD and control children has not been compared directly and the reliability of these measures across different types of movies or movie presentations has not been tested.

In the current study we presented ASD and control children with three short movies, each of which was presented twice. Two of the movies were animated and one was a naturalistic home video, all contained social interactions between at least two individuals. We used both an ROI based and a data driven approach to quantify gaze behavior abnormalities in each of the ASD children. This experimental design enabled us to compare findings across movies, presentations, and eye-tracking measures as well as quantify the gaze pattern idiosyncrasy of individual children. This is particularly important given the large heterogeneity in gaze behaviors of ASD children that is commonly reported in eye tracking studies (Campbell, Shic, Macari, & Chawarska, 2014; Moore et al., 2018; Pierce et al., 2016; Rice et al., 2012).

## Methods

### Participants

Our final analyzed sample included 111 children who were recruited at the Negev Autism Center in Israel (Meiri et al., 2017). This sample consisted of 71 children diagnosed with ASD according to DSM-5 criteria (mean age: 5.1 years old ±1.8, 59 males), and 40 typically developing children (mean age: 4.5 years old ±2.1, 25 males). There were no significant differences in age across the two groups (t(71.4) = 1.5, p = 0.14)). We also performed all of the analyses with a subsample of 28 ASD and 28 control children who were tightly matched for gender (21 males in each group) and age (+/− 2 months). ASD severity was assessed using the Autism Diagnostic Observation Schedule (Lord et al., 2000), and cognitive scores were measured using the Wechsler Preschool and Primary Scale of Intelligence (WPPSI) (Wechsler, 2002) or the Bayley Cognitive Scales (Albers & Grieve, 2007). Children with ASD had a mean ADOS score of 15.4 ±6.2 (range 5-27) and a mean cognitive score of 81.4 ±16 (range 50-117). Parents of all control children completed the Social Responsiveness Scale (SRS) (Rutter, LeCouteur, & Lord, 2015) to ensure that SRS scores were below clinical concern cut-offs (i.e., maximum of 75) (Moody et al., 2017). Control children had a mean SRS score of 35.7 ±14.5 (range 5-67). The final sample described above excluded one control child with an SRS score > 80 as well as 8 ASD children and 2 control children with partial data acquisition (see criteria below). The study was approved by the Soroka Medical Center Helsinki committee and the Ben Gurion University internal review board committee. Written informed consent was obtained from all parents.

### Eye tracking

Children were seated on either an adapted car seat with straps or a comfortable chair (depending on their physical size) such that their head was approximately 60 cm from the screen (head distance was monitored continuously by the eye tracker). The screen was mounted on an adjustable arm from the wall such that children could rest their heads on the back of the seat/chair and minimize head movements. Left eye gaze position was recorded from all participants at a sampling rate of 500 Hz using an EyeLink 1000+ head-free eye tracking system (SR Research Inc. Canada). The eye tracker’s infrared camera was located below the display screen and focused on the eyes of the child. A head tracking sticker was placed on the forehead of the child and the eye tracker was calibrated by presenting five brief salient stimuli at the center and four points of the screen. Calibration was then validated (i.e., stimuli were presented again) to ensure that gaze accuracy was within 2 degrees of initial calibration. Data was acquired and analyzed using Experiment Builder and Data viewer (SR research Inc. Canada). Additional analyses were performed using Matlab (Mathworks Inc., USA).

### Experimental design

After calibrating the eye tracker, children freely viewed 3 short movies, which were presented twice. Each movie was 1.5 minutes long such that the total experiment lasted approximately 10 minutes. Between the movies, a sequence of 10 salient targets was presented to assess the oculomotor control of the children (data not presented). In addition, the accuracy of calibration was tested before and after each movie using a single target, and re-calibration was performed when the error exceeded 2 degrees. Each movie was presented with its original soundtrack through hidden speakers.

The first movie was a segment of the Pixar animation “Jack-Jack Attack”. The segment depicts the adventures of a babysitter who is trying to take care of an infant with supernatural powers. The second movie was a segment of the Walt Disney animation “The Jungle Book”. In the chosen segment Mogli meets the Monkey king who sings and dances while interacting with other monkeys. The third movie was a naturalistic un-cut home-video of a social interaction between two sisters (2 and 5 years old) in a typical messy room with everyday objects.

### Pre-processing and data quality

Children were included in the final sample if their gaze position was recorded successfully for at least 60% of the experiment. We identified segments of lost gaze position or eye blinks lasting <200 ms and linearly interpolated the data to keep the total number of samples identical across subjects. Larger segments of missing data due to movements of the children were labeled and excluded from the analyses.

### Data-analysis

Head and eye regions of interest (ROIs) were manually traced in each frame of the naturalistic movie only. We quantified the percent of time that each child fixated within each ROI relative to the total time that the child watched the movie. Next, we performed three different analyses to assess the differences in moment-by-moment gaze patterns across groups. First, we computed the mean gaze position across all control subjects for each movie frame in each of the three movies. We then computed the Euclidian distance between the gaze position of each subject and the mean gaze position of the control group. Distances were averaged across frames to create a mean distance measure for each child in each movie. When calculating this measure for each of the control children, the mean of the control group was re-calculated without the data of the examined child to ensure statistical independence. Second, we computed the inter-subject correlation of gaze patterns across pairs of children in each group. Correlations were computed separately for the x and y gaze positions and then averaged. We computed the mean inter-subject correlation for each child by averaging across all pairs involving the child. Finally, we computed within-subject correlations by computing the correlation in gaze position across the two presentations of each movie.

A receiver operating characteristic (ROC) analysis was performed by calculating the true positive rate (TPR) and false positive rate (FPR) of identifying ASD children when using different criterion values for each of the eye tracking measures. ROC curves were plotted and the area under the curve (AUC) was computed for each measure. The optimal criterion value for each ROC plot was computed using the Youden Index(Youden, 2006), which is the criterion yielding the largest sum of sensitivity (i.e., TPR) and specificity (i.e., 1 - FPR).

### Statistics

Between group differences were assessed using two sample t-tests with unequal variance, testing for the null hypothesis that the data comes from independent random samples from normal distributions with equal means but not assuming equal variances. Correlations between eye tracking and behavioral measures were all calculated using the Pearson’s linear correlation coefficient.

## Results

### Time gazing at faces and eyes

We manually delineated face and eyes ROIs on each frame of the naturalistic movie. ASD children gazed at the face ROI significantly less than controls (t(82.8) = −2.7, p = 0.01, Figure 1). There was a similar, non-significant, trend for the eyes ROI (t(63.2) = −1.66, p = 0.1). Note the considerable heterogeneity across individuals of both groups and considerable overlap across groups. Performing the same analysis between the age and gender matched groups did not reveal significant differences in the gaze time towards the face (t(53.6) = −0.25, p = 0.8) or eyes (t(52.2) = −0.2, p = 0.85).

**Figure 1.**
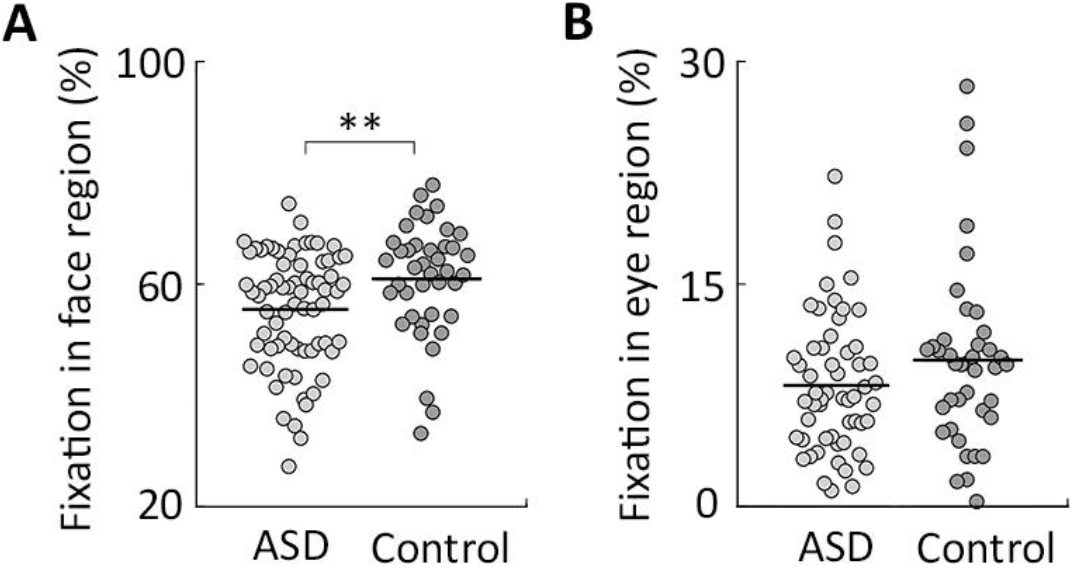
Reduced gaze preference for face and eye ROIs in ASD children. Bee-swarm plots of the relative percentage of time that individual children gazed at the face (A) and eyes (B) ROIs. Light gray: ASD children, Dark gray: Control children. Each circle represents a single child. Asterisks: significant difference across groups (** p=0.01, two sample t-test with unequal variance).

### Gaze distance from the control group mean

We computed the distance between the gaze position of each child and the mean gaze position of the control group for each of the movies (see Methods). This measure quantified the difference between the gaze pattern of each child and the mean gaze pattern of the control group (Figure 2). To ensure statistical independence, when computing this measure for each of the control children, the control group gaze pattern was re-computed without the data of that child (i.e., in a leave-one-out manner).

**Figure 2.**
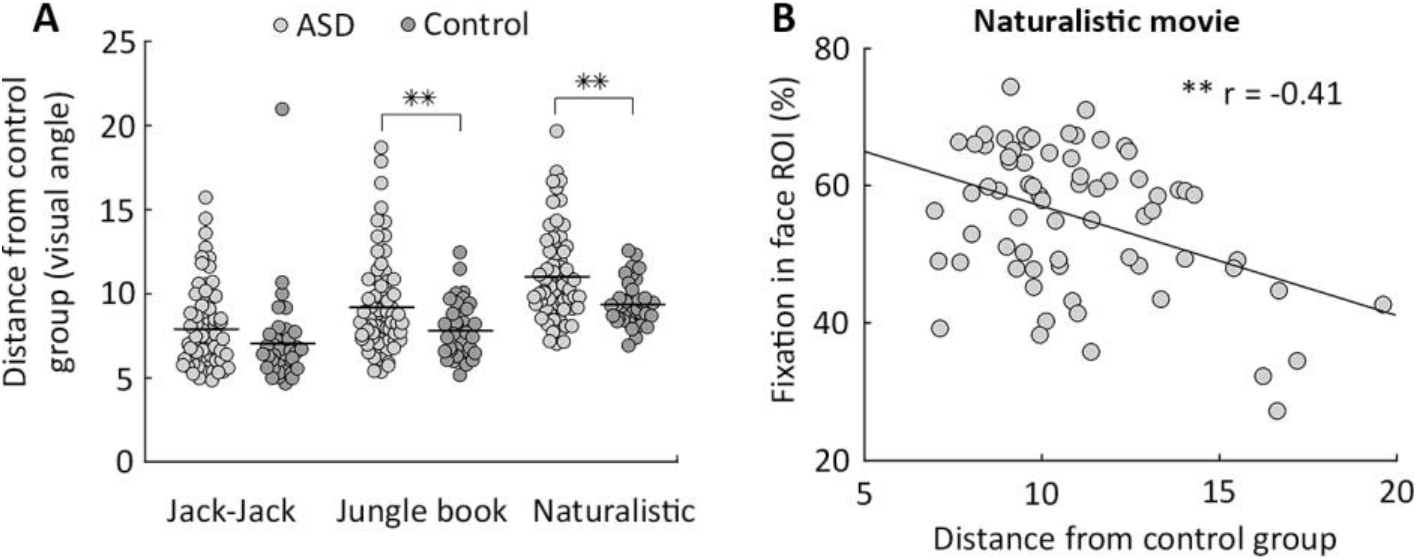
Distance from the mean gaze of the control group. **A.** Bee-swarm plots of individual distances (in degrees) from the control group gaze pattern (mean distance for each movie). **B**. Scatter plot demonstrating the relationship between fixation in the face ROI and the distance from the control group gaze pattern in the naturalistic movie. Line: least squares linear fit. Light gray: ASD children, Dark gray: Control children, Asterisks: significant difference across groups (** p<0.01, two-sample t-test with unequal variance).

ASD children exhibited significantly larger distances from the typical gaze pattern than control children in the Jungle Book (t(109) = 2.8, p = 0.01, d = 0.56) and naturalistic (t(109) = 3.3, p < 0.001, d = 0.73) movies. A non-significant trend in the same direction was also apparent in the Jack Jack movie (t(75.3) = 1.7, p = 0.1, d = 0.34). Similar results were apparent when analyzing the tightly matched ASD and control groups for the Jungle Book (t(54) = 1.9, p = 0.09), naturalistic (t(49.6) = 1.8, p = 0.002), and Jack Jack (t(36.5) = 1.4, p = 0.07) movies.

Individual distances from the typical gaze pattern were significantly, negatively correlated with the time that individual ASD children gazed at the face ROI (r(71) = −0.41, p < 0.001) Figure 2B). This demonstrated that children with a reduced preference for faces, exhibited larger distances from the control group gaze pattern.

The distance from the control group gaze was a stable characteristic of individual ASD children, regardless of movie type or movie presentation (Figure 3). This was demonstrated by significant positive correlations across all pairs of movies (r > 0.64, p < 0.001) and across all pairs of presentations of the same movie (r > 0.66, p < 0.001). We also found a significant positive correlation between the face ROI measure in the two presentations of the naturalistic movie (r = 0.78, p < 0.001), demonstrating that it is also a reliable individual measure.

**Figure 3.**
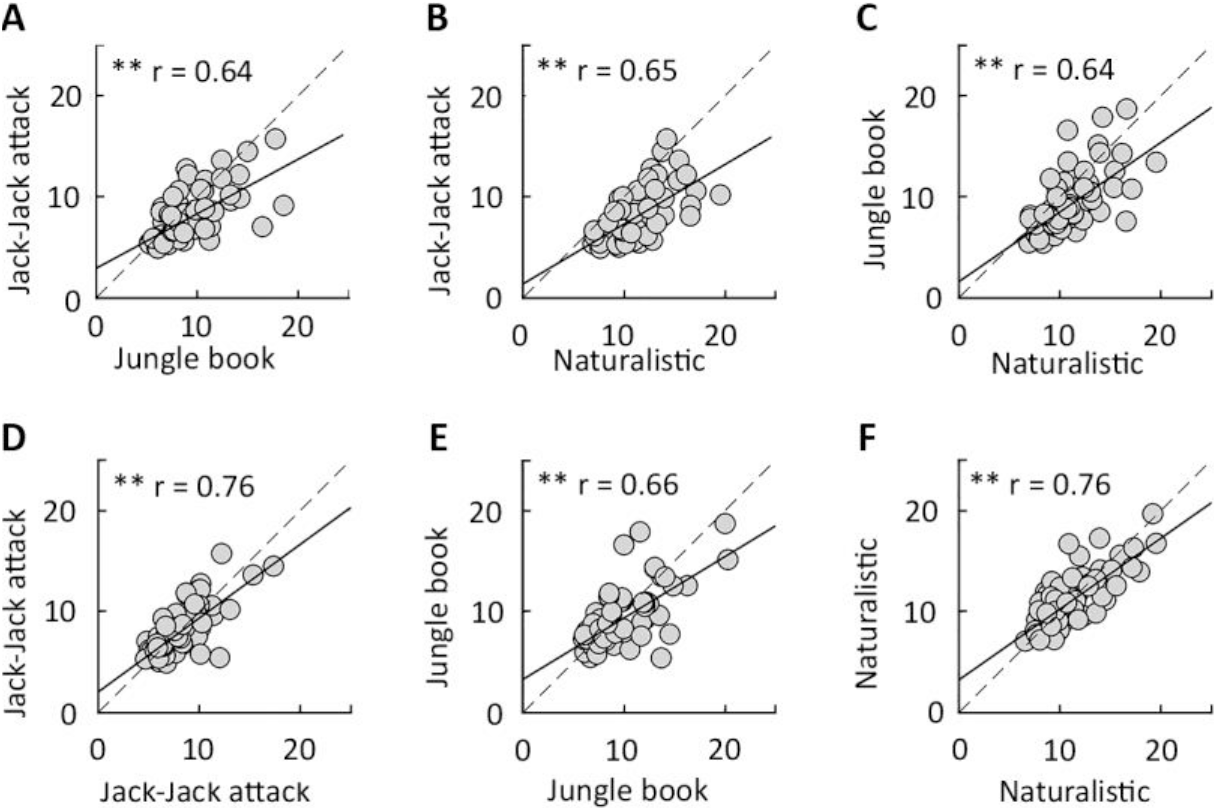
Consistency of distance from control group gaze pattern across movies and presentations. Scatter plots of ASD children demonstrate significant correlations in the distances from control group gaze pattern, across movies (A-C), and across presentations (D-F). Asterisks: significant correlation (** p < 0.001).

To directly compare the utility of the face ROI and gaze distance measures in identifying ASD children, we performed receiver operating characteristic (ROC) analyses for each (Figure 4A). The distance measure exhibited a larger area under the curve (AUC = 0.71, Youden index = 1.4) than the face ROI (AUC = 0.66, Youden index = 1.28) or eyes ROI (AUC = 0.59, Youden index = 1.25) measures. In addition, the optimal cutoff value of the distance measure, calculated using the Youden index, achieved the highest sensitivity (0.72) and specificity (0.68) compared to face ROI (sensitivity = 0.7, specificity = 0.58) or eyes ROI (sensitivity = 0.65, specificity = 0.6). This analysis was performed only for the naturalistic movie where we manually defined the faces and eyes ROIs.

**Figure 4.**
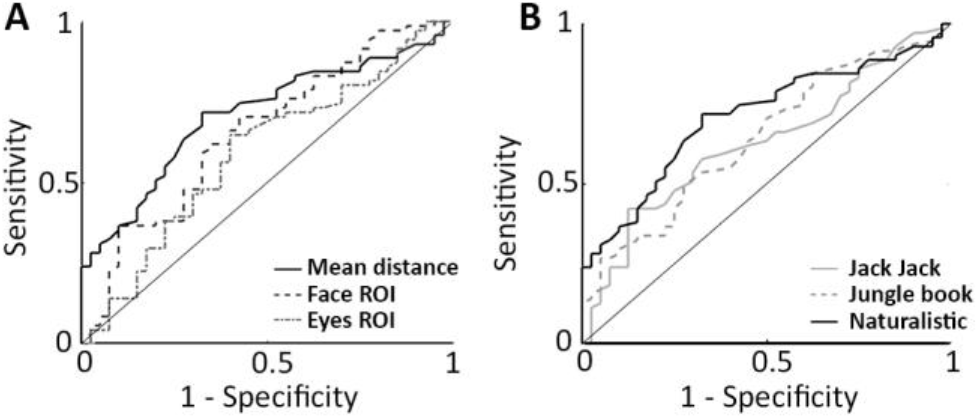
Group separation with ROC analyses. **A:** ROC curves demonstrate the sensitivity and specificity of accurately identifying ASD children based on gaze distance (solid black), face ROI (dashed black), or eyes ROI (dotted gray) measures from the naturalistic movie. **B:** Comparison of ROC curves when using the gaze distance measure in the naturalistic (solid black), Jack-Jack (solid gray), or Jungle Book (dashed gray) movies.

We also performed analogous ROC analyses to compare the utility of the distance measure across the three different movies. This revealed that the best separation between groups was apparent when using the naturalistic movie (AUC = 0.71), followed by the Jungle Book movie (AUC = 0.64), and then the Jack-Jack movie (AUC = 0.63, Figure 4B).

### Inter subject correlations

In a complementary analysis we measured the similarity of gaze patterns across individuals by computing their inter-subject correlations. We computed the correlation in gaze positions throughout each movie across pairs of children from each group. We then computed the average correlation of each child with all others (in their group) to yield an individual measure of inter-subject correlation. Children with ASD exhibited significantly weaker inter-subject correlations with their peers (i.e. more idiosyncratic gaze patterns across individuals) in comparison to controls (Figure 5A) in the naturalistic (t(92.8) = −3.3, p = 0.002), Jack-Jack (t(94.2) = −5.4, p < 0.001), and Jungle book (t(98.2) = −7.9, p < 0.001) movies. Note that different movies elicited different magnitudes of inter-subject correlations due to differences in content and structure. Significant differences across ASD and control groups, however, were apparent regardless of the movie. Hence, the gaze patterns of children with ASD differ from each other more than the gaze patterns of control children.

**Figure 5.**
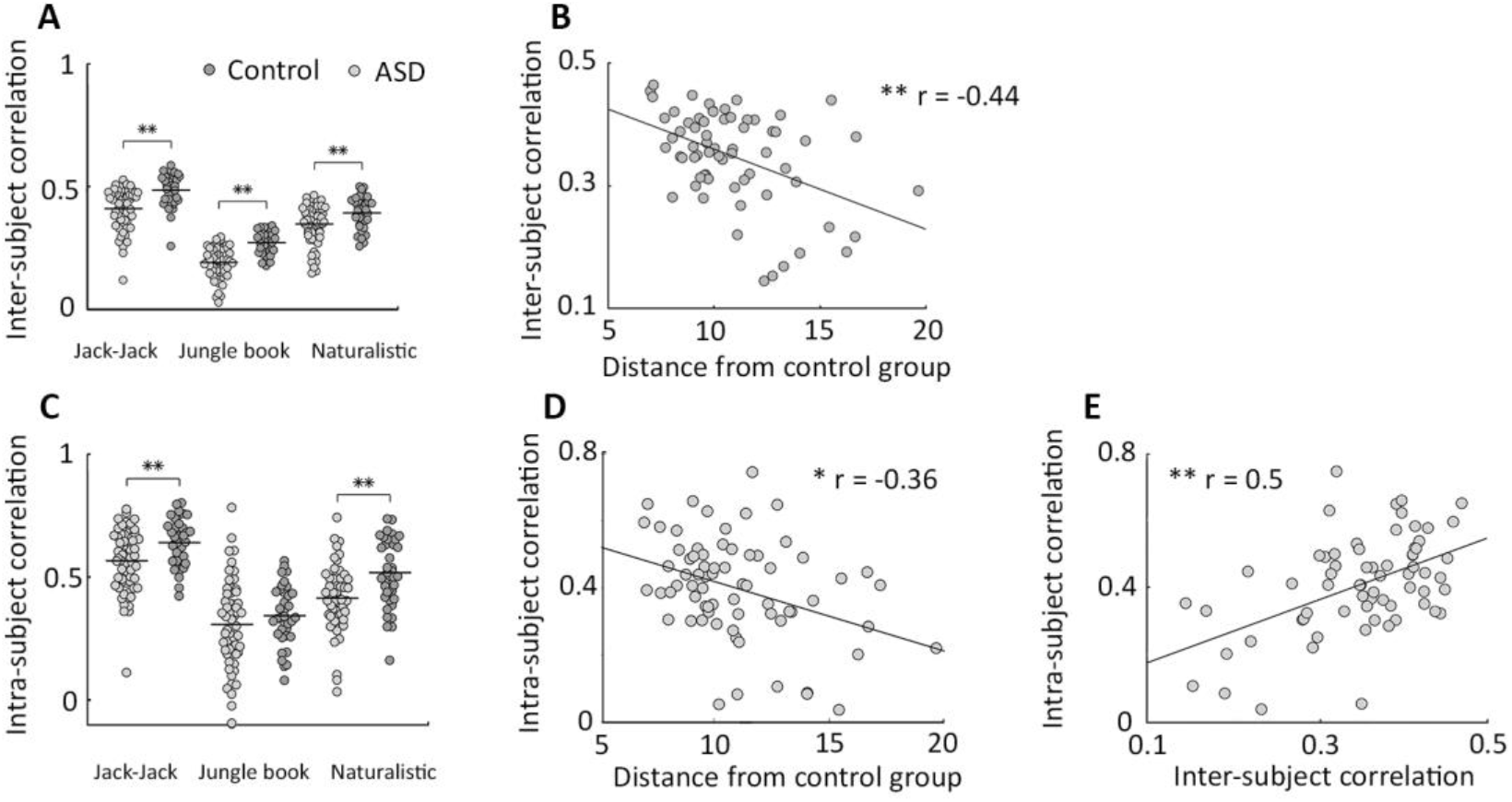
Lower inter- and intra-subject correlations in ASD children. **A.** Bee-swarm plots demonstrating the inter-subject correlation values of individual ASD (light gray) and control (dark gray) children. Horizontal line: group mean. Each circle represents a single child. Asterisks: significant group differences (** p < 0.01, two-sample t-test with unequal variance). **B**. Scatter plot presenting the relationship between inter-subject correlation and distance from the control group gaze. Line: least squares linear fit. Asterisks: significant Pearson’s correlation (* p < 0.05, ** p < 0.01). **C.** Bee-swarm plots demonstrating the intra-subject correlation values of individual ASD and control children (same format as panel A). **D**. Scatter plot presenting the relationship between intra-subject correlation and distance from the control group gaze. **E**. Scatter plot presenting the relationship between intra-subject correlation and inter-subject correlation (panels D and E are in the same format as panel B).

Equivalent results were found when examining the tightly matched ASD and control groups for the naturalistic (t(43.5) = −3.4, p = 0.02), Jack-Jack (t(44.8) = −4.4, p = 0.002), and Jungle book (t(48.6) = −2.4, p < 0.001) movies. ASD children who exhibited lower inter-subject correlations also exhibited larger distances from the control group gaze pattern as demonstrated by a significant negative correlation between the two measures in the naturalistic movie (r(71) = −0.44, p < 0.001) (Fig. 5B).

### Intra subject correlations

To assess within-subject reproducibility in gaze behavior we computed the correlation of gaze patterns across separate presentations of the same movie. ASD children exhibited smaller intra-subject correlations than control children, demonstrating that they viewed each of the movies in a more idiosyncratic and less reproducible manner (Figure 5C). Significant differences were found in the naturalistic (t(84.4) = −3.6, p = 0.001) and Jack-Jack (t(93.2) = 3.4, p = 0.001) movies, and a similar non-significant trend was also apparent in the Jungle book movie (t(91.1) = −1.2, p = 0.2). Note that here too, there were differences in the magnitudes of correlations across the different movies, yet the differences across groups were mostly consistent across all three. Children with lower intra-subject correlations, also exhibited larger distances from the control group gaze pattern as demonstrated by a significant negative correlation across the two measures in the naturalistic movie (Figure 5D, r(71) = −0.36, p < 0.01). The same children also exhibited lower inter-subject correlations as demonstrated by a significant positive correlation across the two measures in the naturalistic movie (Figure 5E, r(71) = 0.5, p < 0.001).

### ASD severity

Significant positive correlations were apparent between the distance from the control group gaze and the total ADOS scores in the naturalistic movie (r(71) = 0.37, p = 0.002) and a similar trend was evident in the jack-jack movie (r(71) = 0.22, p = 0.1). Similarly, significant negative correlations were found between the inter-subject correlation values and the total ADOS scores in the naturalistic (r(71) = −0.46, p < 0.001) and jack-jack (r(71) = −0.52, p < 0.001) movies. Significant negative correlations were also found between the intra-subject correlation values and the total ADOS scores in the naturalistic (r(71) = −0.32, p = 0.01) and jack-jack (r(71) = −0.39, p < 0.001) movies. In contrast, there were no significant correlations between the face or eyes ROI measures and the total ADOS scores in the naturalistic movie (r > −0.12, p > 0.36).

## Discussion

Our results are in agreement with two previous studies (Nakano et al., 2010; Q. Wang et al., 2018) in demonstrating that young children with ASD observe social interactions in a considerably more variable and idiosyncratic manner than control children. This was apparent across several complementary analyses of the children’s gaze patterns. First, the gaze patterns of ASD children deviated more from the mean gaze pattern of the control group in comparison to the gaze patterns of control children (Figure 2). Second, the gaze patterns of the ASD children differed from each other, in comparison to those of control children, as demonstrated by weaker inter-subject correlations in the ASD group (Figure 5A). Third, ASD children exhibited weaker reproducibility in their gaze patterns, as demonstrated by weaker intra-subject correlations, when observing the same movie repeatedly (Figure 5C). All three measures of gaze idiosyncrasy were significantly correlated with each other (Figure 5) and individual differences in all three measures were significantly correlated with ASD symptom severity as assessed by the ADOS. Furthermore, individual magnitudes of gaze idiosyncrasy were reliable individual characteristics, which were significantly correlated across different movies and movie presentations (Figure 3).

Taken together, these results demonstrate that ASD children with more severe symptoms exhibit larger gaze idiosyncrasy, which can be measured reliably using different movies and across different movie presentations. With that said, the largest differences across ASD and control groups were apparent when using the naturalistic home-made video containing a social interaction between two sisters. This suggests that abnormal, idiosyncratic gaze patterns were most pronounced when ASD children observed real-life un-cut interactions of other children (i.e., peers). Indeed, an ROC analysis demonstrated that the naturalistic movie enabled the best separation between ASD and control children, yielding the largest AUC and the best sensitivity and specificity of all three movies (Figure 4B).

In line with other previous studies (Papagiannopoulou et al., 2014), differences across ASD and control groups were also apparent in traditional analyses, which quantified the total time that children gazed at a manually defined face ROI (Figure 1). However, these differences across groups were weaker than those apparent when using gaze idiosyncrasy measures. An ROC analysis demonstrated that the gaze idiosyncrasy measure enabled better separation between the two groups in comparison to the face ROI measure (Figure 4A). Furthermore, ASD symptom severity was not correlated with the face ROI gaze measure.

We, therefore, suggest that utilizing naturalistic movies of social interactions among peers along with measures of gaze idiosyncrasy, offer a potent and reliable technique for quantifying individual gaze abnormalities in ASD children, which are indicative of the children’s ASD symptom severity. This data-driven approach utilizes the temporal complexity of social interactions and offers stronger separation capabilities than traditional analysis techniques using summary statistics of gaze time towards pre-defined ROIs.

### The importance of gaze behavior in social development

Gaze behavior is a remarkably important skill that governs the exposure that an individual has to specific visual information (Henderson, 2003; Schroeder et al., 2010), and simultaneously, conveys potent social information regarding the individual’s state, interests, and intentions (Argyle & Cook, 1976; Emery, 2000). The importance of specific gaze behaviors for early development is apparent in the early emergence of gaze preferences in typically developing infants and toddlers. These include a preference for faces (Baron-Cohen et al., 1997; Batki et al., 2000; Ekman & Friesen, 1971; Frank et al., 2009; Johnson, 2005), biological motion (Fox & McDaniel, 1982; Simion et al., 2008), following the gaze of others (related to joint attention)(Argyle & Cook, 1976; Emery, 2000), and observing the targets of others actions (related to theory of mind)(Flanagan & Johansson, 2003; Oniski & Baillargeon, 2005). These and additional factors, such as visual saliency (Helo et al., 2017), create correlation in the gaze patterns of typically developing toddlers (Franchak et al., 2016) and adults (Shepherd et al., 2010; H. Wang et al., 2012) as they observe movies containing social interactions.

A prominent theory of autism is that abnormalities in early gaze behaviors may explain why children with ASD develop difficulties with social interactions (Klin et al., 2015). The hypothesis is that weaker gaze preferences to social stimuli (Bedford et al., 2012; Chawarska et al., 2012; Chita-Tegmark, 2016; Constantino et al., 2017; Falck-Ytter et al., 2013; W Jones et al., 2008; Warren Jones & Klin, 2013; Klin et al., 2009; Papagiannopoulou et al., 2014; Riby & Hancock, 2009; Rice et al., 2012; Q. Wang et al., 2018) create a situation where children with ASD lack exposure to socially important information and develop gaze behaviors that are less appropriate for social interactions. Interestingly, the lack of visual input and gaze behavior in children who are congenitally blind creates an analogous delay in the development of social skills and these children often fulfill criteria for ASD during childhood, before many of them learn to compensate using other senses (Peter Hobson & Lee, 2010).

An important question is whether children with ASD develop alternative gaze preferences instead of a preference for social stimuli. For example, it has been suggested that individuals with ASD are more attracted to low-level visual saliency (S. Wang et al., 2015). Our results and those of two previous studies (Nakano et al., 2010; Q. Wang et al., 2018) suggest that rather than displaying consistent alternative visual preferences, children with ASD develop variable and idiosyncratic gaze behaviors that are inconsistent across individuals and even within individual across movie presentations.

### Different approaches to measuring gaze behavior

Previous eye tracking studies have used heterogeneous stimuli, measures, and analyses to quantify differences in gaze behavior between ASD and control children. Many have simulated a dyadic interaction with the viewer, where an adult speaks to the viewer in an attempt to capture their attention as a parent may interact with a child (Campbell et al., 2014; Chawarska et al., 2012; Warren Jones & Klin, 2013; Katarzyna, Fred, & Ami, 2010). Other examples include a movie where a child performs yoga-like exercises while facing the camera (Pierce et al., 2016), or movie clips of two children interacting (Nakano et al., 2010). In some cases, the movies were cut such that they included transitions across multiple scenes or clips, while in other cases they contained a single scene (i.e., more naturalistic). These differences in stimulus content, context, and structure are likely to generate differences in the gaze behavior of the viewers (Hasson et al., 2008) as, indeed, apparent in the gaze measures of both groups, which differed across movies in our study (Figures 2A, 5A, and 5C). These differences across stimuli had a clear effect on our ability to separate ASD and control children using gaze patterns from the different movies (Figure 4).

Differences across studies are further compounded by the different measures and analyses used to quantify gaze abnormalities in ASD. While many have used traditional ROI analyses that measure the overall amount of time that children gaze at a particular part or side of the screen (Chawarska et al., 2012; W Jones et al., 2008; Warren Jones & Klin, 2013; Klin et al., 2009; Pierce et al., 2016), more recent studies have compared the moment-by-moment gaze patterns of children (Constantino et al., 2017; Nakano et al., 2010; Q. Wang et al., 2018), which take into account the temporal structure of their gaze behavior rather than focusing only on overall attraction to a single visual feature. Analyzing the rich spatiotemporal dynamics of gaze behavior during the observation of real-life social stimuli is likely to reveal important differences in the gaze behavior of ASD children that may not be captured by previous ROI analyses. Indeed, we found that gaze pattern analyses enabled better separation of ASD and control children than ROI analyses (Figure 4). Furthermore, gaze pattern measures were more strongly correlated with symptom severity as assessed by the ADOS.

A critical focus of future ASD eye tracking studies should be to compare multiple stimuli, measures, and analysis approaches within the same children to demonstrate the benefits of different choices. In addition, assessing the reliability of measures across different movies and across multiple movie presentations, as performed here (Figure 3), is also crucial for developing optimal eye tracking protocols that can identify and quantify symptom severity in children with ASD.

### Heterogeneity of gaze behavior in ASD

ASD is an extremely heterogeneous disorder in terms of its underlying genetics, neurophysiology, and behavioral symptoms (Happé, Ronald, & Plomin, 2006; Jeste & Geschwind, 2014). This heterogeneity is also clearly apparent in the gaze behavior of individual children with ASD. While some have severe gaze abnormalities, others are indistinguishable from typically developing children (Campbell et al., 2014; Pierce et al., 2016; Q. Wang et al., 2018). This was also clearly apparent in our results, where some children with ASD exhibited remarkably idiosyncratic gaze patterns while others did not (Figures 2&5). This heterogeneity in gaze behavior may hold important information, not only for assessing symptom severity, but also for identifying specific endophenotypes within the ASD population (Moore et al., 2018; Rice et al., 2012). We speculate that different stimuli may have different utility for identifying and quantifying specific social ASD symptoms. This specificity may be important for tracking the improvement/deterioration of ASD children over time and for assessing their response to different treatments. It may also enable clinically useful sub-grouping of children with ASD for optimal interventions.

## Conclusions

Eye tracking is likely to be one of the first technologies that will be incorporated into clinical use for assessment of ASD symptoms. Optimizing the ability of eye tracking protocols to identify and quantify specific ASD symptoms will require comparison across the different stimuli, measures, and analysis techniques that have been reported in the literature over the last decade. The current study takes a first important step in this direction by comparing different movies and measures within the same group of ASD and control children. The results indicate that using naturalistic un-cut movies of children’s social interactions along with measures of gaze pattern idiosyncrasy yield the best discrimination between ASD children and controls. Furthermore, these measures were reproducible across movie presentations, demonstrating their reliability, and indicative of individual symptom severity as assessed by the ADOS. Taken together, these results highlight the utility of gaze pattern idiosyncrasy as a reliable eye tracking measure with great potential for clinical utility.

## Acknowledgments

The authors declare no conflict of interest.

This work was funded by ISF grant 961/14 to ID

## Bibliography

Albers, C. A., & Grieve, A. J. (2007). Test Review: Bayley, N. (2006). Bayley Scales of Infant and Toddler Development– Third Edition. San Antonio, TX: Harcourt Assessment. Journal of Psychoeducational Assessment, 25(2), 180–190. https://doi.org/10.1177/0734282906297199

Argyle, M., & Cook, M. (1976). Gaze and mutual gaze. Gaze and mutual gaze. Oxford, England: Cambridge U Press.

Baron-Cohen, S., Wheelwright, S., & Jolliffe, T. (1997). Is there a “language of the eyes”? Evidence from normal adults, and adults with autism or Asperger Syndrome. Visual Cognition, 4(3), 311–331. https://doi.org/10.1080/713756761

Batki, A., Baron-Cohen, S., Wheelwright, S., Connellan, J., & Ahluwalia, J. (2000). Is there an innate gaze module? Evidence from human neonates. Infant Behavior and Development, 23(2), 223–229. https://doi.org/10.1016/S0163-6383(01)00037-6

Bedford, R., Elsabbagh, M., Gliga, T., Pickles, A., Senju, A., Charman, T., & Johnson, M. H. (2012). Precursors to social and communication difficulties in infants at-risk for autism: Gaze following and attentional engagement. Journal of Autism and Developmental Disorders, 42(10), 2208–2218. https://doi.org/10.1007/s10803-012-1450-y

Campbell, D. J., Shic, F., Macari, S., & Chawarska, K. (2014). Gaze response to dyadic bids at 2 years related to outcomes at 3 years in autism spectrum disorders: A subtyping analysis. Journal of Autism and Developmental Disorders, 44(2), 431–442. https://doi.org/10.1007/s10803-013-1885-9

Chawarska, K., Macari, S., & Shic, F. (2012, August). Context modulates attention to social scenes in toddlers with autism. Journal of Child Psychology and Psychiatry, and Allied Disciplines. https://doi.org/10.1111/j.1469-7610.2012.02538.x

Chita-Tegmark, M. (2016). Attention Allocation in ASD: a Review and Meta-analysis of Eye-Tracking Studies. Review Journal of Autism and Developmental Disorders, 3(3), 209–223. https://doi.org/10.1007/s40489-016-0077-x

Constantino, J. N., Kennon-McGill, S., Weichselbaum, C., Marrus, N., Haider, A., Glowinski, A. L., … Jones, W. (2017). Infant viewing of social scenes is under genetic control and is atypical in autism. Nature, 547(7663), 340–344. https://doi.org/10.1038/nature22999

Ekman, P., & Friesen, W. V. (1971). Constants across cultures in the face and emotion. Journal of Personality and Social Psychology, 17(2), 124–129. https://doi.org/10.1037/h0030377

Emery, N. J. (2000). The eyes have it: the neuroethology, function and evolution of social gaze. Neuroscience and Biobehavioral Reviews, 24(6), 581–604.

Falck-Ytter, T., Rehnberg, E., & Bölte, S. (2013). Lack of Visual Orienting to Biological Motion and Audiovisual Synchrony in 3-Year-Olds with Autism. PLoS ONE, 8(7), e68816. https://doi.org/10.1371/journal.pone.0068816

Flanagan, J. R., & Johansson, R. S. (2003). Action plans used in action observation. Nature, 424(6950), 769–771. https://doi.org/10.1038/nature01861

Fox, R., & McDaniel, C. (1982). The perception of biological motion by human infants. Science, 218(4571), 486–487. https://doi.org/10.1126/science.7123249

Franchak, J. M., Heeger, D. J., Hasson, U., & Adolph, K. E. (2016). Free Viewing Gaze Behavior in Infants and Adults. Infancy, 21(3), 262–287. https://doi.org/10.1111/infa.12119

Frank, M. C., Vul, E., & Johnson, S. P. (2009). Development of infants’ attention to faces during the first year. Cognition, 110(2), 160–170. https://doi.org/10.1016/j.cognition.2008.11.010

Frazier, T. W., Klingemier, E. W., Parikh, S., Speer, L., Strauss, M. S., Eng, C., … Youngstrom, E. A. (2018). Development and Validation of Objective and Quantitative Eye Tracking−Based Measures of Autism Risk and Symptom Levels. Journal of the American Academy of Child and Adolescent Psychiatry, 57(11), 858–866. https://doi.org/10.1016/j.jaac.2018.06.023

Happé, F., Ronald, A., & Plomin, R. (2006). Time to give up on a single explanation for autism. Nature Neuroscience, 9(10), 1218–1220. https://doi.org/10.1038/nn1770

Hasson, U., Landesman, O., Knappmeyer, B., Vallines, I., Rubin, N., & Heeger, D. J. (2008). Neurocinematics: The Neuroscience of Film. Projections, 2(1), 1–26. https://doi.org/10.3167/proj.2008.020102

Helo, A., van Ommen, S., Pannasch, S., Danteny-Dordoigne, L., & Rämä, P. (2017). Influence of semantic consistency and perceptual features on visual attention during scene viewing in toddlers. Infant Behavior and Development, 49, 248–266. https://doi.org/10.1016/j.infbeh.2017.09.008

Henderson, J. M. (2003). Human gaze control during real-world scene perception. Trends in Cognitive Sciences. https://doi.org/10.1016/j.tics.2003.09.006

Hensch, T. K. (2005). Critical period plasticity in local cortical circuits. Nature Reviews. Neuroscience, 6(11), 877–888. https://doi.org/10.1038/nrn1787

Jeste, S. S., & Geschwind, D. H. (2014). Disentangling the heterogeneity of autism spectrum disorder through genetic findings. Nature Reviews. Neurology, 10(2), 74–81. https://doi.org/10.1038/nrneurol.2013.278

Johnson, M. H. (2005, October 1). Subcortical face processing. Nature Reviews Neuroscience. Nature Publishing Group. https://doi.org/10.1038/nrn1766

Jones, W, Carr, K., & Klin, A. (2008). Absence of preferential looking to the eyes of approaching adults predicts level of social disability in 2-year-old toddlers with autism spectrum disorder. Archives of General Psychiatry, 65(8), 946–954. https://doi.org/10.1001/archpsyc.65.8.946

Jones, Warren, & Klin, A. (2013). Attention to eyes is present but in decline in 2-6-month-old infants later diagnosed with autism. Nature, 504(7480), 427–431. https://doi.org/10.1038/nature12715

Katarzyna, C., Fred, V., & Ami, K. (2010). Limited Attentional Bias for Faces in Toddlers With Autism Spectrum Disorders. Archives of General Psychiatry, 67(2), 178. https://doi.org/10.1001/archgenpsychiatry.2009.194

Klin, A., Lin, D. J., Gorrindo, P., Ramsay, G., & Jones, W. (2009). Two-year-olds with autism orient to non-social contingencies rather than biological motion. Nature, 459(7244), 257–261. https://doi.org/10.1038/nature07868

Klin, A., Shultz, S., & Jones, W. (2015, March 1). Social visual engagement in infants and toddlers with autism: Early developmental transitions and a model of pathogenesis. Neuroscience and Biobehavioral Reviews. Pergamon. https://doi.org/10.1016/j.neubiorev.2014.10.006

Lord, C., Risi, S., Lambrecht, L., Cook, E. H. J., Leventhal, B. L., DiLavore, P. C., … Rutter, M. (2000). The Autism Diagnostic Schedule – Generic: A standard measures of social and communication deficits associated with the spectrum of autism. Journal of Autism and Developmental Disorders, 30(3), 205–223. https://doi.org/10.1023/A:1005592401947

Meiri, G., Dinstein, I., Michaelowski, A., Flusser, H., Ilan, M., Faroy, M., … Menashe, I. (2017). Brief Report: The Negev Hospital-University-Based (HUB) Autism Database. Journal of Autism and Developmental Disorders, 47(9), 2918–2926. https://doi.org/10.1007/s10803-017-3207-0

Moody, E. J., Reyes, N., Ledbetter, C., Wiggins, L., DiGuiseppi, C., Alexander, A., … Rosenberg, S. A. (2017). Screening for Autism with the SRS and SCQ: Variations across Demographic, Developmental and Behavioral Factors in Preschool Children. Journal of Autism and Developmental Disorders, 47(11), 3550–3561. https://doi.org/10.1007/s10803-017-3255-5

Moore, A., Wozniak, M., Yousef, A., Barnes, C. C., Cha, D., Courchesne, E., & Pierce, K. (2018). The geometric preference subtype in ASD: Identifying a consistent, early-emerging phenomenon through eye tracking. Molecular Autism, 9(1), 19. https://doi.org/10.1186/s13229-018-0202-z

Nakano, T., Tanaka, K., Endo, Y., Yamane, Y., Yamamoto, T., Nakano, Y., … Kitazawa, S. (2010). Atypical gaze patterns in children and adults with autism spectrum disorders dissociated from developmental changes in gaze behaviour. In Proceedings of the Royal Society B: Biological Sciences. https://doi.org/10.1098/rspb.2010.0587

Oniski, K. K., & Baillargeon, R. (2005). Do 15-month-old infants understand false beliefs? Science, 308(5719), 255–258. https://doi.org/10.1126/science.1107621

Papagiannopoulou, E. A., Chitty, K. M., Hermens, D. F., Hickie, I. B., & Lagopoulos, J. (2014). A systematic review and meta-analysis of eye-tracking studies in children with autism spectrum disorders. Social Neuroscience, 9(6), 610–632. https://doi.org/10.1080/17470919.2014.934966

Peter Hobson, R., & Lee, A. (2010). Reversible autism among congenitally blind children? A controlled follow-up study. Journal of Child Psychology and Psychiatry, 51(11), 1235–1241. https://doi.org/10.1111/j.1469-7610.2010.02274.x

Pierce, K., Marinero, S., Hazin, R., McKenna, B., Barnes, C. C., & Malige, A. (2016). Eye tracking reveals abnormal visual preference for geometric images as an early biomarker of an autism spectrum disorder subtype associated with increased symptom severity. Biological Psychiatry, 79(8), 657–666. https://doi.org/10.1016/j.biopsych.2015.03.032

Riby, D., & Hancock, P. (2009). Looking at movies and cartoons: Eye-tracking evidence from Williams syndrome and autism. Journal of Intellectual Disability Research, 53(2), 169–181. https://doi.org/10.1111/j.1365-2788.2008.01142.x

Rice, K., Moriuchi, J. M., Jones, W., & Klin, A. (2012). Parsing heterogeneity in autism spectrum disorders: Visual scanning of dynamic social scenes in school-aged children. Journal of the American Academy of Child and Adolescent Psychiatry, 51(3), 238–248. https://doi.org/10.1016/j.jaac.2011.12.017

Rutter, M., LeCouteur, A., & Lord, C. (2015). Autism Diagnostic Interview - Revised (ADI-R). Statewide Agricultural Land Use Baseline 2015, 1. https://doi.org/10.1017/CBO9781107415324.004

Sasson, N. J., & Elison, J. T. (2012). Eye Tracking Young Children with Autism. Journal of Visualized Experiments, (61). https://doi.org/10.3791/3675

Schroeder, C. E., Wilson, D. A., Radman, T., Scharfman, H., & Lakatos, P. (2010, April). Dynamics of Active Sensing and perceptual selection. Current Opinion in Neurobiology. https://doi.org/10.1016/j.conb.2010.02.010

Senju, A., & Johnson, M. H. (2009). Atypical eye contact in autism: Models, mechanisms and development. Neuroscience & Biobehavioral Reviews, 33(8), 1204–1214. https://doi.org/10.1016/j.neubiorev.2009.06.001

Shepherd, S. V, Steckenfinger, S. A., Hasson, U., & Ghazanfar, A. A. (2010). Human-Monkey Gaze Correlations Reveal Convergent and Divergent Patterns of Movie Viewing. Current Biology, 20(7), 649–656. https://doi.org/10.1016/j.cub.2010.02.032

Simion, F., Regolin, L., & Bulf, H. (2008). A predisposition for biological motion in the newborn baby. Proceedings of the National Academy of Sciences, 105(2), 809–813. https://doi.org/10.1073/pnas.0707021105

Tanaka, J. W., & Sung, A. (2016). The “Eye Avoidance” Hypothesis of Autism Face Processing. Journal of Autism and Developmental Disorders, 46(5), 1538–1552. https://doi.org/10.1007/s10803-013-1976-7

Wang, H., Freeman, J., Merriam, E., Hasson, U., & Heeger, D. (2012). Temporal eye movement strategies during naturalistic viewing. Journal of Vision, 12(1), 16–16. https://doi.org/10.1167/12.1.16

Wang, Q., Campbell, D. J., Macari, S. L., Chawarska, K., & Shic, F. (2018). Operationalizing atypical gaze in toddlers with autism spectrum disorders: A cohesion-based approach. Molecular Autism, 9(1), 25. https://doi.org/10.1186/s13229-018-0211-y

Wang, S., Jiang, M., Duchesne, X. M., Laugeson, E. A., Kennedy, D. P., Adolphs, R., & Zhao, Q. (2015). Atypical Visual Saliency in Autism Spectrum Disorder Quantified through Model-Based Eye Tracking. Neuron, 88(3), 604–616. https://doi.org/10.1016/j.neuron.2015.09.042

Wechsler, D. (2002). Wechsler Preschool and Primary Scale of Intelligence. Wechsler Preschool and Primary Scale of Intelligence, 120–130. https://doi.org/10.1007/978-1-4419-1698-3_866

Youden, W. J. (2006). Index for rating diagnostic tests. Cancer, 3(1), 32–35. https://doi.org/10.1002/1097-0142(1950)3:1<32::aid-cncr2820030106>3.0.co;2-3

